# Biochemical Functions of the Membrane-Binding Domain of CARMIL

**DOI:** 10.64898/2026.01.05.697744

**Authors:** Olivia L. Mooren, Patrick McConnell, James D. DeBrecht, John A. Cooper

## Abstract

Actin assembly at membranes is often associated with proteins with domains that bind and regulate heterodimeric actin capping protein (CP). CP-binding domains can target CP to the membrane and activate CP by promoting dissociation of its stoichiometric inhibitor V-1. The capping protein binding region (CBR) of CARMIL includes a CPI motif and a CSI motif, followed by a membrane binding (MB) domain. The MB domain is necessary for the function of CARMIL in cells, and it is sufficient for targeting GFP to the plasma membrane of cells. Here, we investigated the mechanism and significance of the relationship of the MB domain to CP activity, including capping of actin filament barbed ends and promotion of Arp2/3-nucleated actin assembly. We found that the MB domain is able to bind to lipid-coated beads, bringing the CPI and CSI motifs to the bead, and thus activating CP to promote Arp2/3-based actin assembly. In addition, we discovered that the MB domain can dissociate from the lipid membrane once CP binds; this observation may help account for the long-standing quandary as to how activated CP is released from the membrane and how CP functions to activate Arp2/3-mediated actin assembly near the membrane. We also report that the MB domain released from the membrane enhances the ability of the CPI and CSI domains to activate CP. Thus, the CARMIL MB domain has multiple biochemical functions regulating actin assembly at a membrane. First, it can target CARMIL, CP, and barbed ends to the plasma membrane. Second, the MB domain can leave the membrane, and this promotes the uncapping of capped barbed ends and activates soluble CP, with greater ability than seen with the membrane-attached state.

## Introduction

Actin filament assembly is necessary for cells to change their shape, move components about the cytoplasm, and migrate from one location to another (1). The location of actin filaments determines when and where myosin-based movements, such as contractions of muscle tissue and platelets, occur. In addition, the polymerization of actin subunits into filaments generates force that drives the movement of membranes, including plasma membranes. These forces change the shape of cells and power their migration.

Regulation of actin filament assembly is provided by proteins that bind directly to actin and by regulatory molecules that bind to actin-binding proteins (2). Assembly of filaments from subunits often occurs at barbed ends (3), which can be bound and controlled by the heterodimeric α/β capping protein (CP) (4). CP is regulated by the protein V-1 (aka myotrophin), which binds directly to CP on the top of its mushroom-shaped structure. The V-1 binding site overlaps with the binding site for the barbed end of the actin filament, sterically blocking the CP / actin interaction. CP is also regulated by proteins with motifs that bind directly to CP at a separate location distinct from the barbed-end binding site (recently reviewed in (5)). These motifs are termed “CP-Interacting (CPI)” motifs (6), and they are found in a wide variety of otherwise unrelated proteins (7).

CARMIL proteins are a family of CP regulators (7, 8). They contain a Capping Protein-Interaction (CPI) motif, a CARMIL-Specific Interacting (CSI) motif (6), and a membrane-binding (MB) basic-hydrophobic (BH) motif. The three domains are found in tandem in the unstructured C-terminal half of vertebrate CARMILs (diagrammed in Figure 1, panel A). CARMILs of invertebrates and protozoa contain the CPI motif but lack the CSI and MB domains (8).

**Figure 1.**
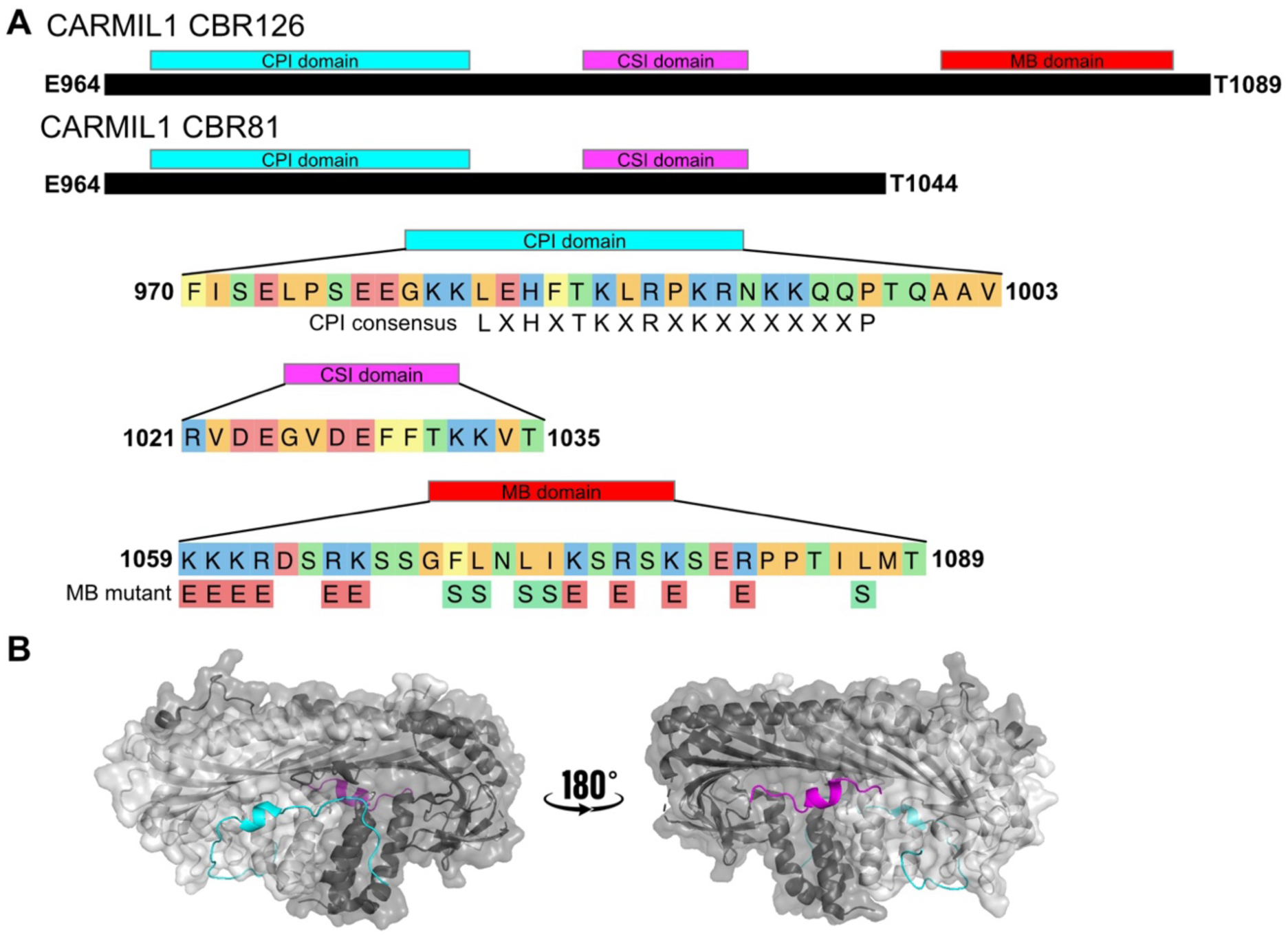
Domain architecture of the CP-binding region (CBR) of human CARMIL1. **A.** The polypeptides in this study are shown, with the three important regions expanded for additional detail. The conserved CPI consensus sequence is listed below the CPI domain. The set of amino-acid mutations used to abrogate membrane binding is shown below the MB domain. **B.** Crystal structures (modified from PDB 3LK3) of CP (alpha subunit in dark grey and beta subunit in light grey) with the CARMIL1 CPI-motif peptide (cyan) and CSI-motif peptide (magenta).

The CPI motif binds directly to the stalk of CP, at a site distinct from the site for binding V-1 and actin. The CSI motif also binds to the stalk of CP, continuing from the CPI-binding site (6) (see Figure 1, panel B). The linker between the CPI and CSI motifs is not involved in binding to or regulation of CP (9). The BH motif of the MB domain was discovered with a sequence search algorithm designed to identify short regions of basic and hydrophobic residues that bind to acidic lipid membranes (10). That study identified mouse CARMIL1, several myosin-I’s, and several PAK kinases (10). A subsequent study showed that the MB domain is present in the conserved sequences of all three vertebrate CARMIL isoforms – 1, 2 and 3 (11). The MB domain of all three CARMILs is sufficient for localizing GFP to the plasma membrane, and the MB domain of human CARMIL2 is necessary for the cellular functions of CARMIL2 in cultured-cell systems of actin-based motility (11).

In a previous study using a biochemical reconstitution system (12), we obtained results in support of a model for CP regulation developed by Fujiwara, Hammer and colleagues (5, 13). One fundamental element of the model is stoichiometric inhibition of CP’s capping activity by V-1, which sterically blocks CP’s binding site for the barbed end of the actin filament. A second essential feature of the model is allosteric regulation induced by CARMIL binding to the CP / V-1 complex.

The CPI and CSI motifs bind to CP at a distant site, away from the site that binds actin and V-1. CARMIL binding to CP induces a conformational change that weakens V-1 binding by promoting its dissociation from the CP / V-1 complex (14, 15). The net effect of CARMIL binding is to counteract the inhibitory action of V-1 and thereby activate CP. Active CP is one of the components essential for actin assembly mediated by activation of Arp2/3 complex (16). Thus, this model provides a novel mechanism for regulation of actin assembly by any mechanism that creates free barbed ends, including nucleation by formins, EnaVASP, and SPIN90 proteins (17–19). Ultimately, CARMIL regulates CP’s capping activity because the conformational change induced by CARMIL CBR binding to CP affects both the actin-binding site of CP and the V-1-binding site of CP, which overlap with each other (5, 6, 14, 15, 20–23).

Our previous study included a fragment of the CP-binding region (CBR) of human CARMIL1 termed “CBR115”, which contains the CPI and CSI motifs. CARMILs appear to function at membranes, and the MB domain with its BH motif is always in close proximity directly C-terminal to the CPI and CSI motifs. We asked how the MB domain and its binding to lipids affect the regulation of CP, using a single CARMIL CBR fragment encompassing all three motifs. We examined how the MB domain affects the binding of CP to actin using functional assays for actin assembly, and conversely, how the interaction with CP affects the ability of a CARMIL fragment to bind to lipids.

We found that the MB domain is sufficient for targeting the CARMIL fragment to an acidic phospholipid membrane, and that the targeted CARMIL fragment is able to promote Arp2/3-mediated actin assembly by promoting V-1 dissociation from CP / V-1 complex. Conversely, we found that binding of CP to the CARMIL CBR fragment leads to partial dissociation of the CBR fragment from the lipid. This unexpected observation provides insight into how CARMILs might leave a membrane, and it provides for dissociation of CP from actin filament barbed ends, as suggested by studies of CP / actin association lifetimes in cells (24, 25).

## Results

The CARMIL family of proteins is distinct among vertebrate CPI-motif proteins because they contain two conserved regions following the CPI domain – the CSI domain and the MB domain (Figure 1, panel A). The CPI and CSI motifs were defined by structural analysis showing their direct interactions with CP, their sequence conservation, and their biochemical and cellular effects on CP function (5, 6, 9, 23). We investigated the biochemical functions of the three domains using a fragment of the CP-binding region (CBR) of CARMIL1, termed “CBR126”, which includes all three of the CBR domains. First, we examined how the MB domain affects the capping activity of the CPI / CSI domain construct. Second, we examined how the interaction of the MB domain with lipids affects CP activity, in terms of a) capping actin barbed ends and b) promoting Arp2/3-nucleated actin assembly. For (a), we used traditional pyrene-actin capping assays in solution, and for (b), we employed a bead surface-based actin reconstitution system employed in our recent study (12) with the modification of adding anionic phospholipids to the bead surface to mimic the plasma membrane.

To begin, we confirmed that the MB domain can target CARMIL1-CBR126 to a lipid membrane, as shown in cells (11), using a biochemical assay of liposome sedimentation. CARMIL1-CBR126 sedimented almost completely in the presence of liposomes, and it did not sediment in the absence of liposomes (Figure 2, panel A and B, blue vs magenta). This result confirms the prediction, from studies of GFP-MB fusion proteins expressed in cells (11), that the MB domain is sufficient to achieve targeting to a biochemical lipid membrane.

**Figure 2.**
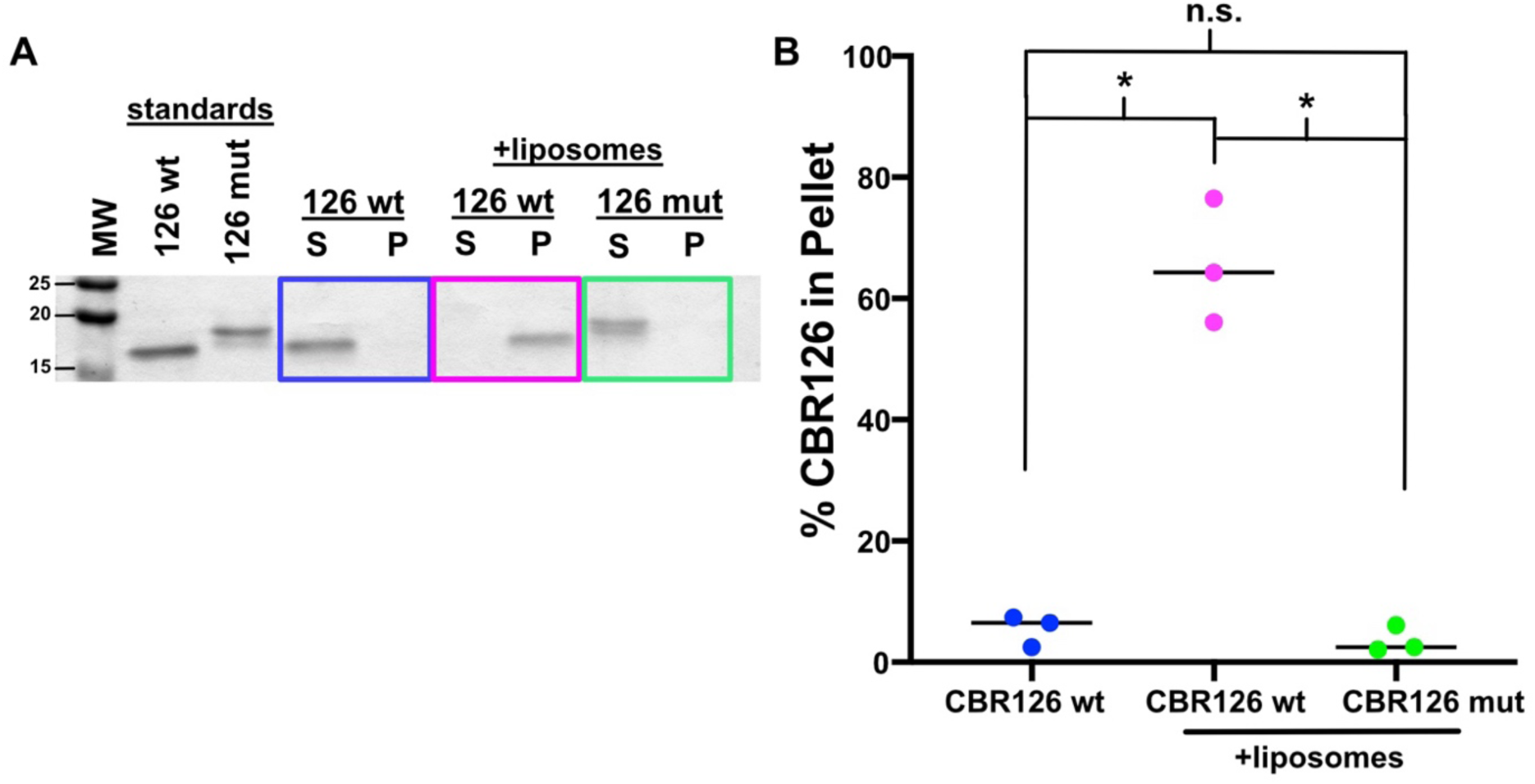
CARMIL1-CBR126 associates with phospholipid membranes via the membrane binding (MB) domain. **A.** SDS-PAGE of protein from the supernatant (S) and pellet (P) following centrifugation in liposome sedimentation assays. CBR126 wt pellets almost completely with liposomes (magenta), whereas the membrane binding mutant (CBR126 mut, green) is primarily in the supernatant. **B.** The percentage of CBR126 in the pellet plotted in panel B is compared to a standard sample loaded on the same gel (see panel A). Each point is a separate experiment with p (*) < 0.01 in a Welch’s t test (26). The horizontal line is the median.

We next asked how the association of CBR126 with a lipid membrane affects its ability to interact with CP and regulate actin assembly. As a negative control for membrane binding of CBR126, we designed and constructed loss-of-function membrane-binding mutants of the MB motif. We changed basic and hydrophobic amino acids known to contribute to the membrane-binding function (basic to acidic, hydrophobic to hydrophilic) (Figure 1, panel A) (10, 11). First, we mutated the 13-amino-acid core of the BH motif in CARMIL1, previously found to be necessary for CARMIL2 membrane targeting (11). This mutation decreased the association of CBR126 with lipids by approximately half in the liposome sedimentation assay (Suppl. Figure 1). We then made a longer mutation of the MB domain, covering a span of 29 amino-acid residues (Figure 1, panel A). This longer mutation completely abolished the interaction of CBR126 with liposomes in the sedimentation assay (Figure 2, panel A and B, magenta versus green). Thus, the residues of the BH motif were confirmed to be responsible for the membrane-binding ability of CBR126. In subsequent experiments testing the functional roles of the MB domain, we compared the activity of CBR126 wt with the longer 29-aa mutant, termed “CBR126 mut.”

In addition, we deleted the entire MB domain by truncating CBR126 between the CSI and MB domains, creating a fragment termed “CBR81”, to address the question of whether the MB domain has a negative or positive effect on the CPI / CSI interaction with CP. CBR81 did not pellet with liposomes in the sedimentation assay (Suppl. Figure 1), again showing the necessity of the MB region for interaction of the CBR region of CARMIL1 with lipids.

### Effect of MB Domain on Actin Capping Activity in Solution

Next, we asked how the MB domain affects the ability of the CP-binding region (CBR) to regulate actin capping activity of CP in solution, using pyrene-actin polymerization assays. Increasing concentrations of CBR126 wt completely inhibited the actin capping activity of CP (Figure 3, panel A, and Suppl. Figure 2 for full titration curves). The CBR126 mutant also inhibited CP well, but not as effectively as the CBR126 wt (Figure 3, panels A vs B, blue curves). The binding affinities of CBR126 wt and CBR126 mut for CP, measured by ITC, were similar (Suppl. Figure 3). We also tested the truncated CARMIL1 CBR81 fragment, which lacks the MB domain. CBR81 was about as effective as CBR126 wt at inhibiting CP in the actin capping assay (Figure 3, blue curves of panels A and C).

**Figure 3.**
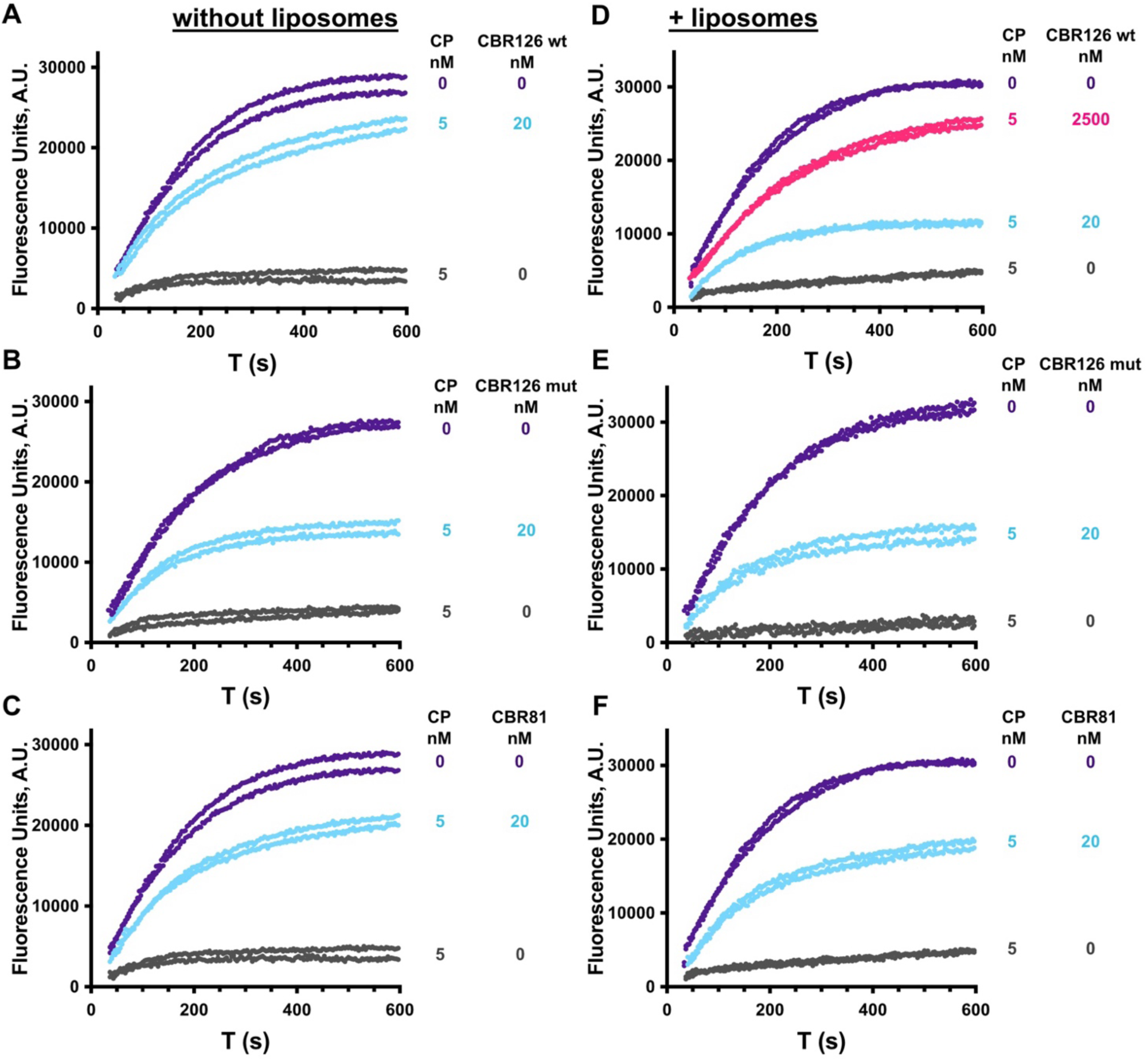
Effects of CARMIL1 CBR126 MB domain and lipids on actin capping activity of CP. Pyrene actin polymerization assays seeded with barbed ends performed in the absence (panels A - C) and presence (panels D – F) of liposomes. Each curve is one independent experiment. Purple curves are absence of CP, and grey curves are with addition of CP. **Panel A.** CBR126 wt inhibits CP (blue curves). **Panels B and C.** Similar to panel A, with CBR126 MB mut (panel B) or CBR81 (panel C). **Panels D – F.** Analogous to Panels A – C, with addition of liposomes. Far more CBR126 wt (125x) is needed to inhibit CP (pink curves in panel D vs blue curves in panel A). **E and F.** Liposomes have no apparent effect on the activity of CBR126 mut or CBR81. NB: These curves are selected from a complete titration set shown in Supplemental Figure 2.

Since the MB domain is always located just C-terminal to the CPI and CSI motifs in CARMILs, we hypothesized that membrane interaction might affect the ability of CPI / CSI to interact with CP. To investigate whether and how lipid binding affects the interaction activity of CBR126 with CP, we added lipid membranes in the form of liposomes to the pyrene-actin-based capping assays. When liposomes were present, 125x more CBR126 wt was required to obtain the same level of CP inhibition as when liposomes were absent from the assay mixture (Figure 3, panel D pink curve compared to panel A blue curve). As a negative control, we tested the CBR126 mut in a dose-response assay (Figure 3 and Suppl. Figure 2); the results confirm that liposomes do not significantly alter the actin capping activity of the MB mutant (Figure 3, panel E compared to panel B). As an additional negative control, we tested the activity of CBR81, truncated to lack the MB domain entirely. We found that CBR81 was also not significantly affected by the presence of liposomes (Figure 3, compare panel F with panel C). Overall, the important conclusion is that liposome membrane interaction decreases the biochemical activity of CBR126 on CP with respect to actin capping.

To further investigate the biochemical interactions of CBR126 with membranes and CP, we added CP to liposome sedimentation assays. As shown above (Figure 2), CBR126 sedimented with liposomes. When an equimolar amount of CP was added to the reaction mixture, the amount of CBR126 sedimenting with the liposomes was less, indicating that the interaction of CBR126 with CP decreases the membrane binding of CBR126 (Suppl. Figure 4, panel A). In a titration experiment, increasing amounts of CP in the mixture led to corresponding decreases in the amount of CBR126 sedimenting with liposomes (Suppl. Figure 4, panel C) along with decreases in the fraction of CP sedimenting with liposomes (Suppl. Figure 4, panel D). CP was present in the pellet fraction along with CBR126 (Suppl. Figure 4, panel B), showing that CP is able to bind to CBR126 that is bound to membrane. Note that CP alone, in the absence of CBR126, did not pellet with liposomes. We conclude that CP can bind to membranes via its interaction with CBR126, and that the CBR126 / CP complex can leave the membrane. In other words, CBR126 can bind to the lipid membrane and to CP. The two binding states are dynamic and interconvert; they are not exclusive of one another.

### Arp2/3-mediated Actin Assembly at Lipid-coated Surface

Previously, we demonstrated that CP can be activated by CARMIL1 at a surface by releasing CP from V-1, using a biomimetic bead assay for Arp2/3-mediated actin assembly (12). In that study, an active fragment of CARMIL1 fused to GST was coupled to glutathione-coupled beads. Here, to create a surface that mimics the membrane bilayer, we deposited anionic phospholipid onto a bead. We documented lipid deposition by fluorescence imaging of beads with a fluorescent lipid tracer and by scanning electron microscopy (Suppl. Figure 5).

First, we asked whether CBR associated with the lipid-coated bead surface is able to activate CP and promote Arp2/3-nucleated actin assembly (12). In our recombinant system Arp2/3 complex is activated by a VVCA fragment of N-WASp. To couple the VVCA to the bead, we used His-tagged VVCA and Ni-functionalized lipids. On addition of Arp2/3 complex and actin, actin polymerization was observed around the bead, in the form of asymmetric tails. The actin assembly required CP (Figure 4, top row), and addition of V-1 suppressed actin tail growth by inactivating CP (Figure 4, 2^nd^ row). The results document that Arp2/3-mediated actin polymerization can occur from the lipid-coated beads, that active CP is necessary, and that V-1 inhibits CP and prevents actin growth.

**Figure 4.**
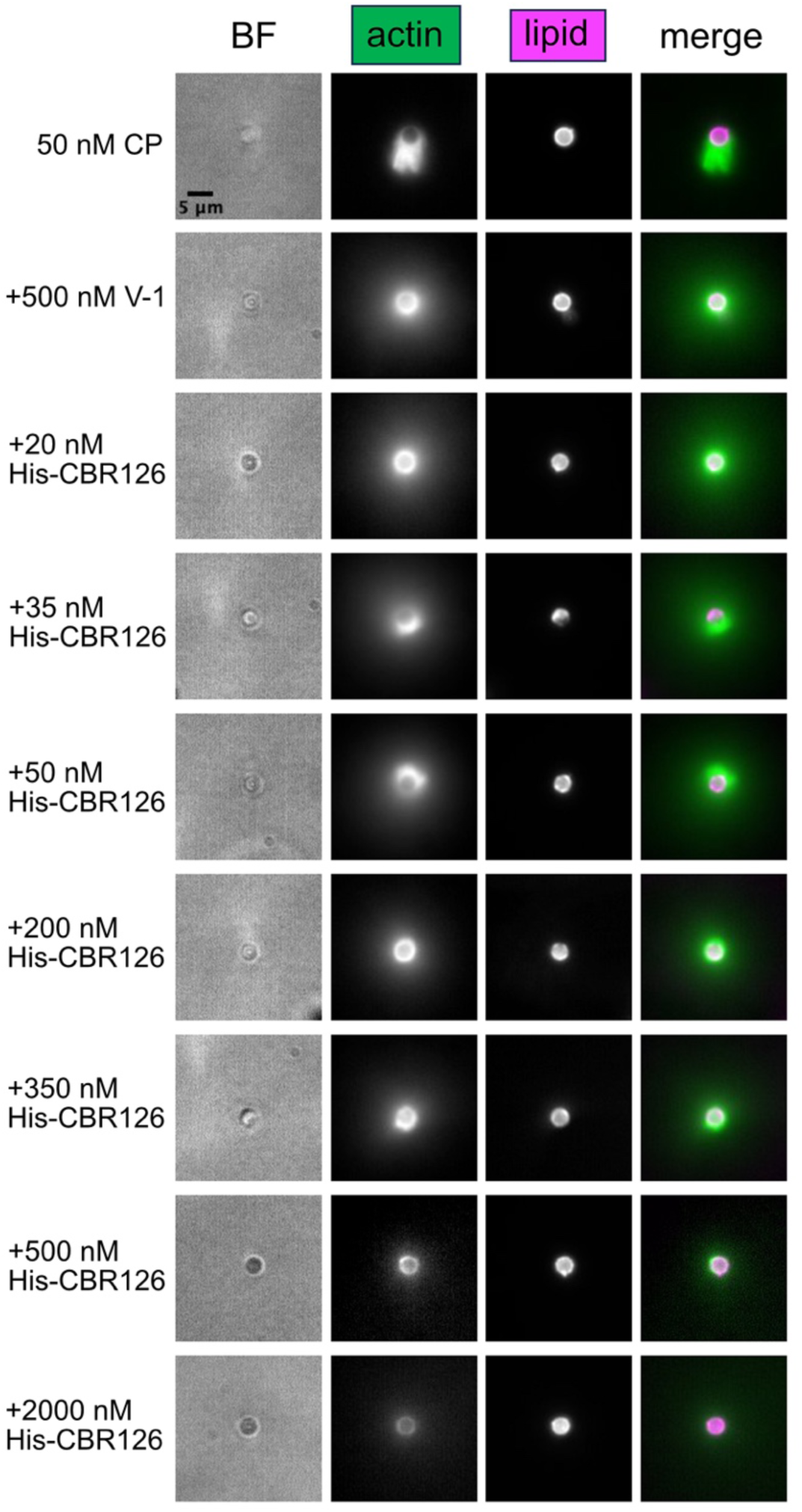
Actin filament network assembly on lipid-coated beads with CP, V-1, and CBR126. Asymmetric tails of actin filament networks generated by incubating Ni-functionalized and fluorescent lipid-coated beads with His-VVCA (N-WASP), followed by addition of 100 nM Arp2/3 complex, 5 µM profilin-actin, and 50 nM CP for 30 min (top row). Addition of 500 nM V-1 to the reaction mixture resulted in F-actin growing from the bead surface as a symmetric ring and a diffuse cloud around the bead (second row). Addition of low concentrations of His-CBR126 resulted in asymmetric F-actin tail growth from the bead (rows labeled 35 nM and 50 nM), and higher concentrations inhibited actin growth (row labeled 2000 nM).

As a control experiment, we compared the results here with CBR126 and lipid-coated beads with those from our previous study (12)., which used a shorter fragment of the CBR (CBR115), lacking part of the MB domain (12) and glutathione beads. For that comparison, we used a His-tagged version of CBR126, designed to couple CBR126 directly to the Ni-functionalized lipid-coated beads, independently of the MB domain. Indeed, the actin network assembly results were similar to those of our previous study. In an experiment titrating the amount of His-CBR126 added to the beads, low amounts of His-CBR126 caused asymmetric actin tail growth from the bead surface, presumably by activating CP from CP/V-1 complex. With higher amounts of His-CBR126, the actin tail growth was inhibited, by the direct effect of CBR126 on the actin capping activity of CP (Figure 4, 20 nM – 2000 nM, rows three thru nine). The inhibition of CP activity in this assay is consistent with previous studies showing CARMIL CBR at high levels lowers the actin capping activity of CP (23, 27).

To assess the contribution of the MB domain to His-CBR126 activity on CP and actin network growth in the bead assay, we attached His-CARMIL1-CBR126 mut to the beads instead of His-CBR126. We did not observe much difference in the asymmetric actin growth from beads compared to His-tagged CBR126 wt at the same concentrations (Figure 5, panel A, compare rows three and four). We conclude that, in the context of the bead assays when CARMIL1 is bound to the bead surface via a His-tag, the CPI and CSI domains are the primary components of the CBR important for influencing CP activity and actin network assembly.

**Figure 5.**
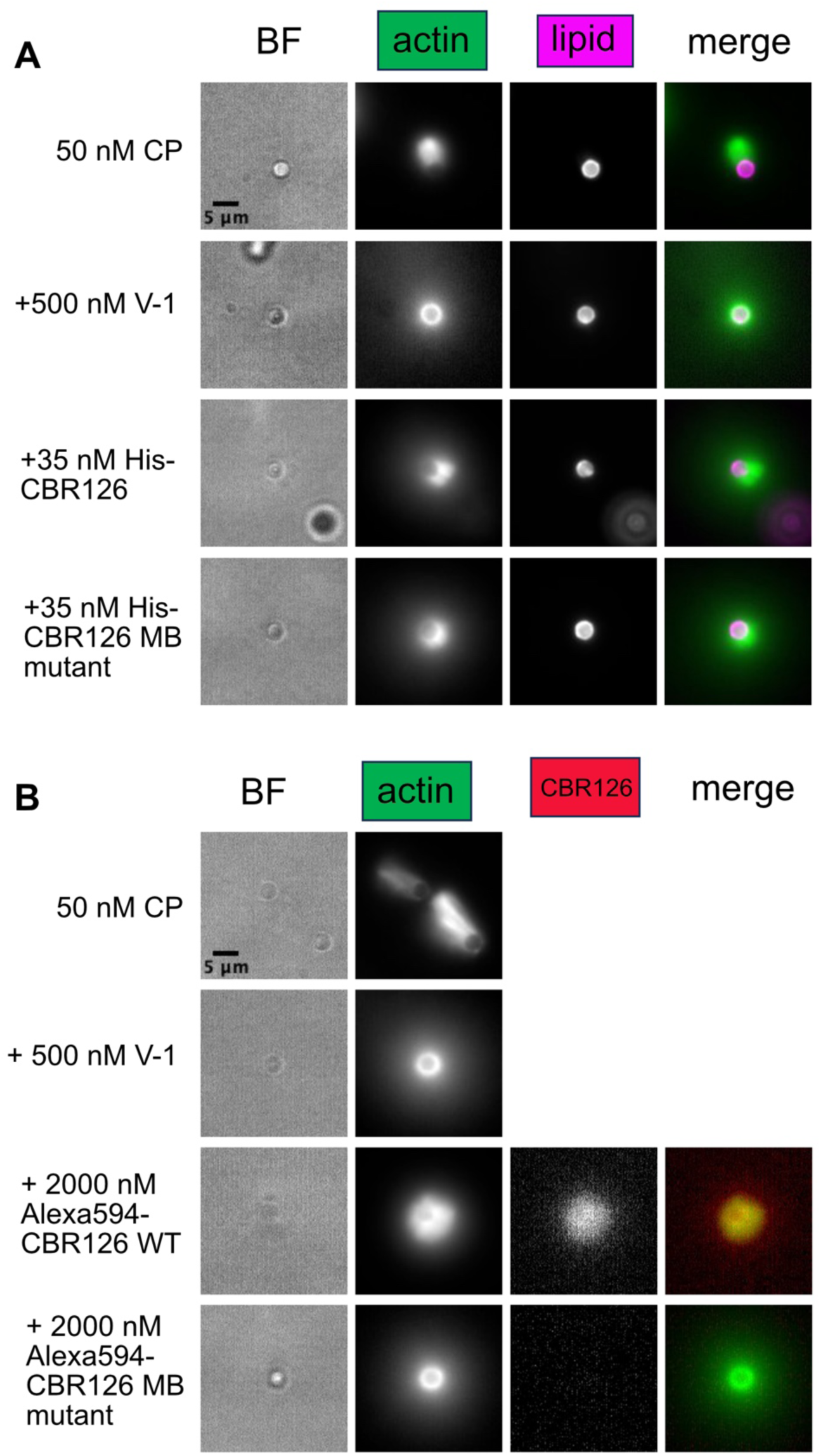
Membrane binding domain (MB) important for function because of CARMIL localization. **A.** His-tagged MB mutants cause actin network to grow asymmetrically from the bead surface in a mixture of 100 nM Arp2/3 complex, 5 µM profilin-actin, 50 nM CP, and 500 nM V-1 (30-minute time points) similar to His-CBR126 wt. **B.** Untethered CBR126 associates with the lipid beads via the MB domain and displays more robust asymmetric actin growth than tethered His-CBR126. MB mutants do not associate with the lipid beads and do not show the same effects on the actin network.

To address further whether dynamic lipid association plays an important role in CBR effect on CP activity and actin network growth, we used CBR126 that did not contain the His-tag to investigate whether the MB domain of the CBR of CARMIL1 is sufficient to target CBR to the lipid-coated beads. We fluorescently labeled the His-tag-less CBR126 and followed its localization with fluorescence imaging. Fluorescent CBR126 appeared on the surface of the lipid-coated bead, prior to adding all the components of the actin assembly mixture (Suppl. Figure 5, panel A), indicating that the MB domain of CARMIL1 is sufficient for localizing CARMIL1 to the bead surface. After adding all the actin assembly components, we observed robust asymmetric actin growth on beads (Figure 5, panel B, row 3), at levels greater than what we observed for His-tagged CBR126. As the actin polymerization reaction progressed over time (30 min), the intensity of CBR126 fluorescence on the bead surface decreased. (Suppl. Figure 6, panel B). For some beads, CBR126 fluorescence was observed to overlap with the fluorescence of the diffuse F-actin cloud network by two-color imaging (Figure 5, panel B, row 3). For other beads, CBR126 fluorescence was not observed. As an additional test of the CBR126 localization, we performed immunofluorescence staining with anti-CBR126. This experiment also showed CBR126 was at times distributed throughout the F-actin network (Suppl. Figure 6, panel C). Parenthetically, we note that greater amounts of the tag-less CBR126 compared to His-tagged-CBR126 are required to produce positive effects on the growth of the actin network. This result is consistent with the pyrene actin polymerization assays in which the activity of CBR126 has less effect on the activity of CP when liposomes are present.

To confirm that the MB domain is responsible for the lipid membrane localization and for the functional effects observed with His-less CBR126, we used a fluorescent derivative of the mutant; CBR126 mut did not appear on the lipid-coated bead surface prior to adding the actin assembly reaction mixture (Suppl. Figure 6, panel A) and no asymmetric actin assembly was observed (Figure 5, panel B, bottom row).

In sum, the results show the following: a) the MB domain of CARMIL1 CBR126 is sufficient for localizing CBR126 to the lipid membrane; b) CBR126 targeted to the bead surface via a lipid membrane is sufficient to activate CP and thus actin assembly; and c) CBR126 can transition off the lipid-coated bead surface when CBR126 binds to CP due to the dynamic nature of the interaction of the BH motif with lipid.

## Discussion

CARMILs bind directly to CP and regulate its ability to bind V-1 and cap actin filaments (5). The CP-binding region (CBR) of CARMILs is an ∼130-aa segment of the intrinsically disordered C-terminal half of the ∼1400-aa polypeptide. The CBR includes three domains in a tandem array (from N- to C-term): the CPI domain, the CSI domain, and the MB domain. The CPI and CSI domains bind directly to the surface of CP, and they induce a conformational change in CP that affects its distant binding sites for F-actin and V-1.

The conserved position of the MB domain within the three-fold tandem array of the CBR raises questions about how the MB domain itself or binding of the MB domain to a lipid membrane might affect the interaction of the CBR with CP. Previous studies demonstrated that the MB domain is necessary for the function of CARMIL2 in human cultured cells (11) and that the MB domain of any one of three vertebrate CARMIL isoforms is found to be sufficient to target intracellular GFP to cell membranes (11).

We asked how the MB domain and lipids affect the ability of the CPI and CSI domains to regulate the biochemical activity of CP in assays of actin capping and Arp2/3-mediated actin nucleation. First, we found that the MB domain is sufficient to target the CBR to a lipid membrane, in binding assays with liposomes or lipid-coated beads. Second, we found that the CBR, when targeted to a lipid-coated bead, is able to activate CP (by inhibition of V-1 binding) and thus promote actin assembly nucleated by Arp2/3 complex. Third, we found that lipid binding decreases the ability of CBR to inhibit the actin capping activity of CP in solution, based on pyrene-actin polymerization assays with liposomes.

However, the binding of the MB domain to a lipid membrane is a dynamic interaction based on set of relatively weak interactions of the basic and hydrophobic residues with the acidic phospholipids of the membrane. We found that CBR would partition between the membrane and the soluble phases, and that the binding of CP would increase the fraction of CBR that partitioned off the membrane into the soluble phase.

Together, these observations imply that CBR that dissociates from the lipid membrane is relatively more effective in binding CP than is membrane-bound CP. Greater binding implies greater activation of CP based on dissociation of CP / V-1 complex and greater inhibition of CP’s capping of the barbed ends of actin filaments. Both of these effects, dissociation of V-1 and inhibition of capping, are expected to promote actin assembly near the membrane.

Indeed, the dynamic nature of the MB domain / membrane interaction was revealed in both our liposome sedimentation experiments and our lipid-coated bead-based actin assembly reconstitution system. First, in liposome sedimentation experiments, a fraction of CP sedimented with liposomes, but only when CBR was present as predicted. CBR with a mutated form of the MB, lacking basic and hydrophobic residues, was not able to bind liposomes and thus did not suffice to recruit CP to the liposomes. More important, when CP was present and bound to CBR, then the fraction of CBR associated with the liposomes was decreased.

Second, in the lipid-coated bead-based actin assembly reconstitution experiments, CBR was initially associated completely with the bead surface. However, as the actin filaments assembled and grew away from the bead surface, the CBR was observed to leave the bead surface and move into the actin filament network that formed a cloud around the bead. In addition, we observed more robust actin assembly with the tag-less CBR126 than with the His-tagged CBR126. This implies that the ability of CBR to leave the membrane can be important for function.

### Model for CARMIL / CP Function at Membranes

Together, these observations substantiate the existence of multiple states for CARMIL and CP that have been proposed and speculated about in studies of actin-based cell motility. One can envision multiple paths between the states in our biochemical results, which would correspond with states of actin filament membrane attachment and actin-based assembly in cells. The paths are diagrammed in the cartoon model of Figure 6.

**Figure 6.**
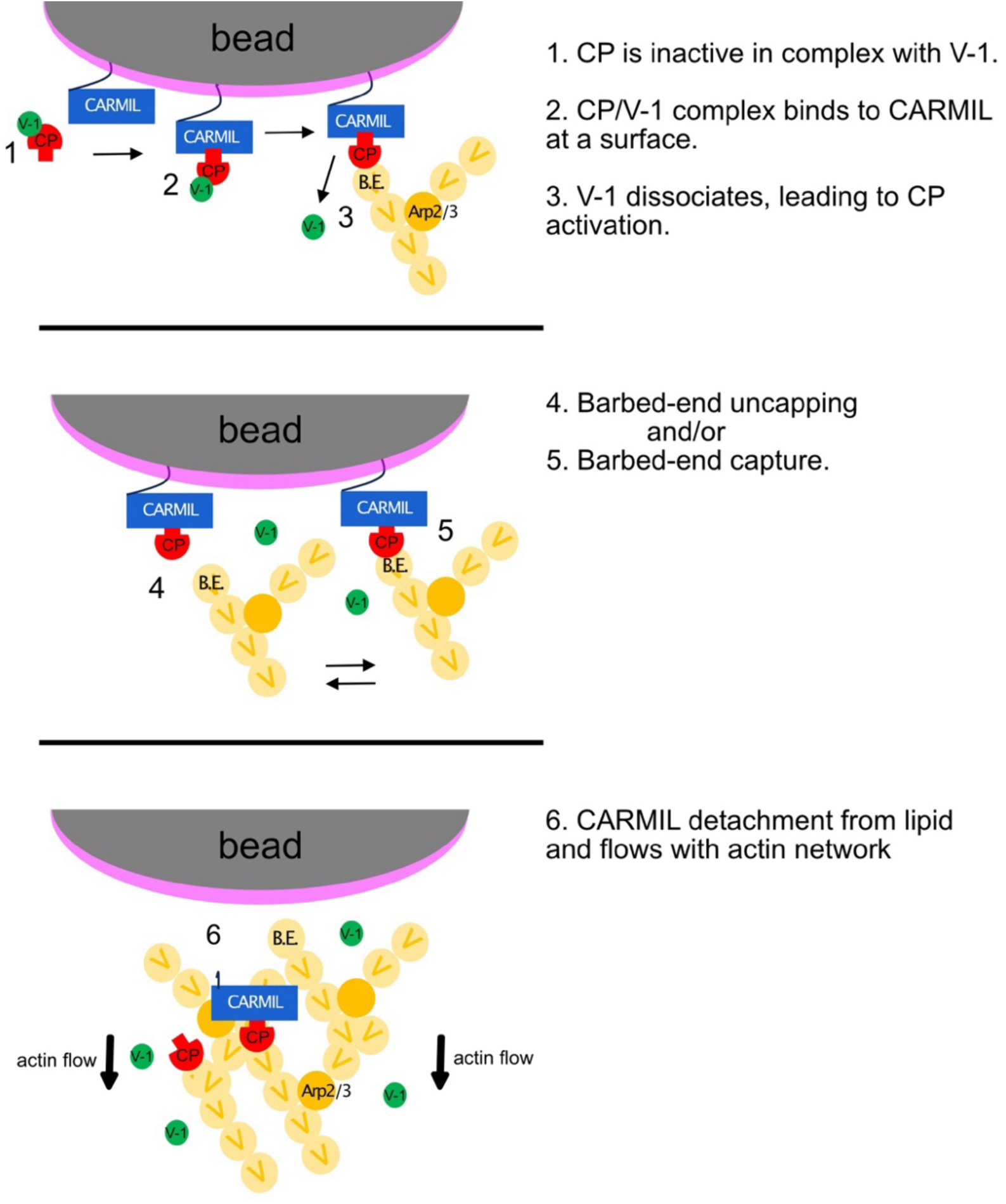
Model of regulatory cycles for CP actin capping. 1. CP bound to V-1 in the cytoplasm is inactive. **2.** CP / V-1 binding to CARMIL promotes V-1 dissociation. **3.** Free CP binds barbed ends and promotes Arp2/3-nucleated polarized actin growth at the bead surface. **4 & 5.** Near the bead surface, CARMIL can a) promote uncapping of a capped barbed end to allow filament growth or b) capture a capped actin filament. Dynamic association of CP with barbed end - “loose / leaky” capper. **6.** Dynamic association of CARMIL with lipid: CARMIL can leave the bead surface and stay bound to CP as the actin filament network grows and flows away from the bead surface.

First, CARMIL, represented by CBR in this analysis, is sufficient to bind CP to a membrane and to activate CP for barbed-end capping (Figure 6). Thus, membrane-bound CARMILs may contribute to the morphology of actin filament barbed ends appearing as attached to or embedded in electron-dense material adjacent to the plasma membrane (28–30). Second, CARMILs can convert capped barbed ends to uncapped ones, as documented with single-molecule biochemical experiments (13). As known from previous studies, active CP near the membrane suffices to promote Arp2/3-based actin assembly from new free barbed ends (31, 32). In addition, the dynamic nature of the MB / membrane interaction implies that CARMIL is able to leave the membrane (Figure 6). Once off the membrane, the ability of the CBR to promote uncapping (dissociation of CP from the barbed end) is increased, as is expected to occur during retraction of a cell process (33, 34).

### Open Questions and Future Directions

First, this study utilized only the CBR fragment of CARMIL, not full-length protein. The field recognizes that the other regions of CARMILs, outside the CBR, have additional biochemical functions related to interactions with other biomolecules and regulation of CARMIL activity. Evidence exists that CARMILs can be autoinhibited and regulated by signaling molecules (5).

An important future direction for the field will be to define the relative abundance of the states considered and detected here, along with the relative timing and rate of transitions of CARMILs and CP among those states.

## Conclusions

The membrane-binding domain of CARMIL is sufficient to attach the CP-binding region to lipids. On the surface of a lipid-coated bead, the CBR attached via its MB domain is able to activate CP and thereby promote Arp2/3 complex mediated actin assembly.

The CBR, when placed on the membrane, via its MB domain, is much less active for CP interaction compared to its activity in solution. When the CBR dissociates from and leaves the membrane, to function in solution, the presence of the MB domain slightly increases the ability of CBR to interact with CP.

The interaction of CARMIL with lipids is dynamic, converting between lipid-bound and free in solution. The transition between on and off the membrane may help account for the attachment of actin filament barbed ends to the membrane, as well as for detachment and uncapping. These transitions and states correspond to ones that occur at the leading edge of motile cells.

## Acknowledgments

We are grateful to Dr. Michael Onken and members of our laboratory for advice and assistance. Supported by NIH grant R35 GM144082 to J.A.C. We acknowledge the Washington University Center for Cellular Imaging (WUCCI) for its electron microscopy studies, which are supported by Washington University School of Medicine, The Children’s Discovery Institute of Washington University and St. Louis Children’s Hospital (CDI-CORE-2015-505 and CDI-CORE-2019-813), and the Foundation for Barnes-Jewish Hospital (3770 and 4642).

## Supplementary Figures

**Suppl. Figure 1.**
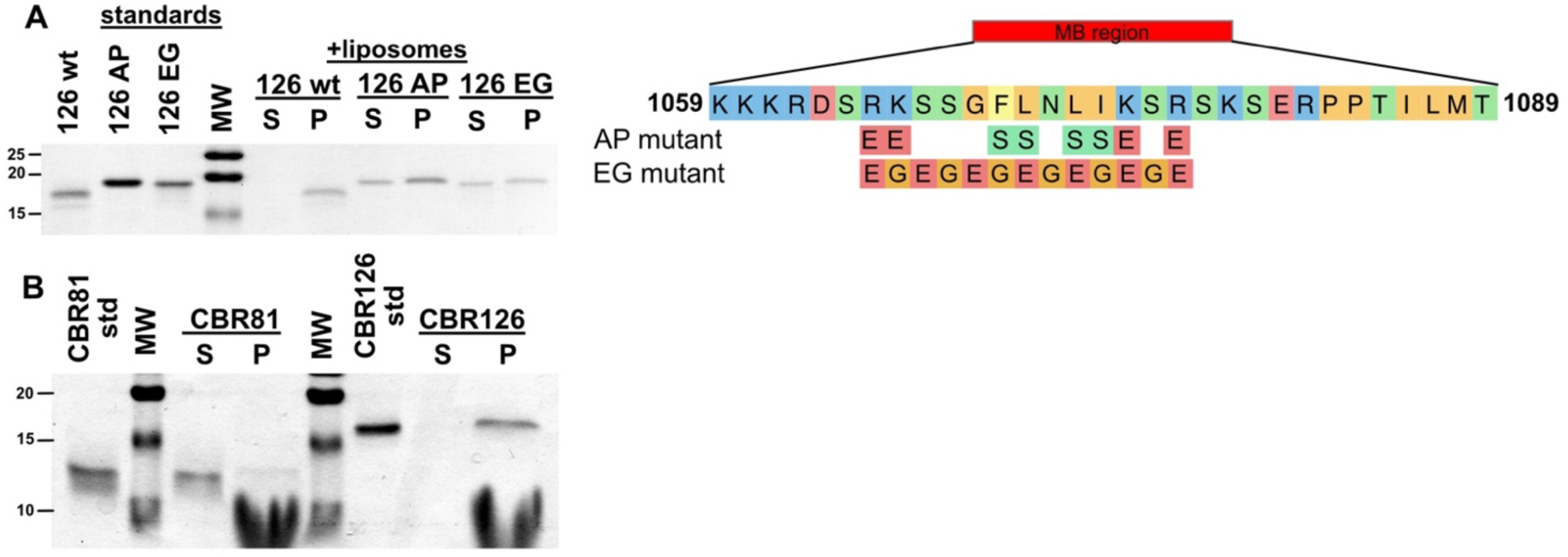
MB domain mutants and membrane binding. **A.** SDS-PAGE of sedimentation assays of MB mutants with two different sets of changes to a core 13-aa sequence, diagrammed on the right. One set, labeled “AP mutant”, reverses basic/hydrophobic residues to acidic and polar residues, and the other set, labeled “EG mutant”, consists of wholesale changes of residues to an alternating pattern of E and G. Each mutant shows decreased but not abrogated binding to liposomes in the sedimentation assay. **B.** SDS-PAGE of sedimentation assay showing that CBR81, which lacks the MB domain, did not associate with liposomes.

**Suppl. Figure 2.**
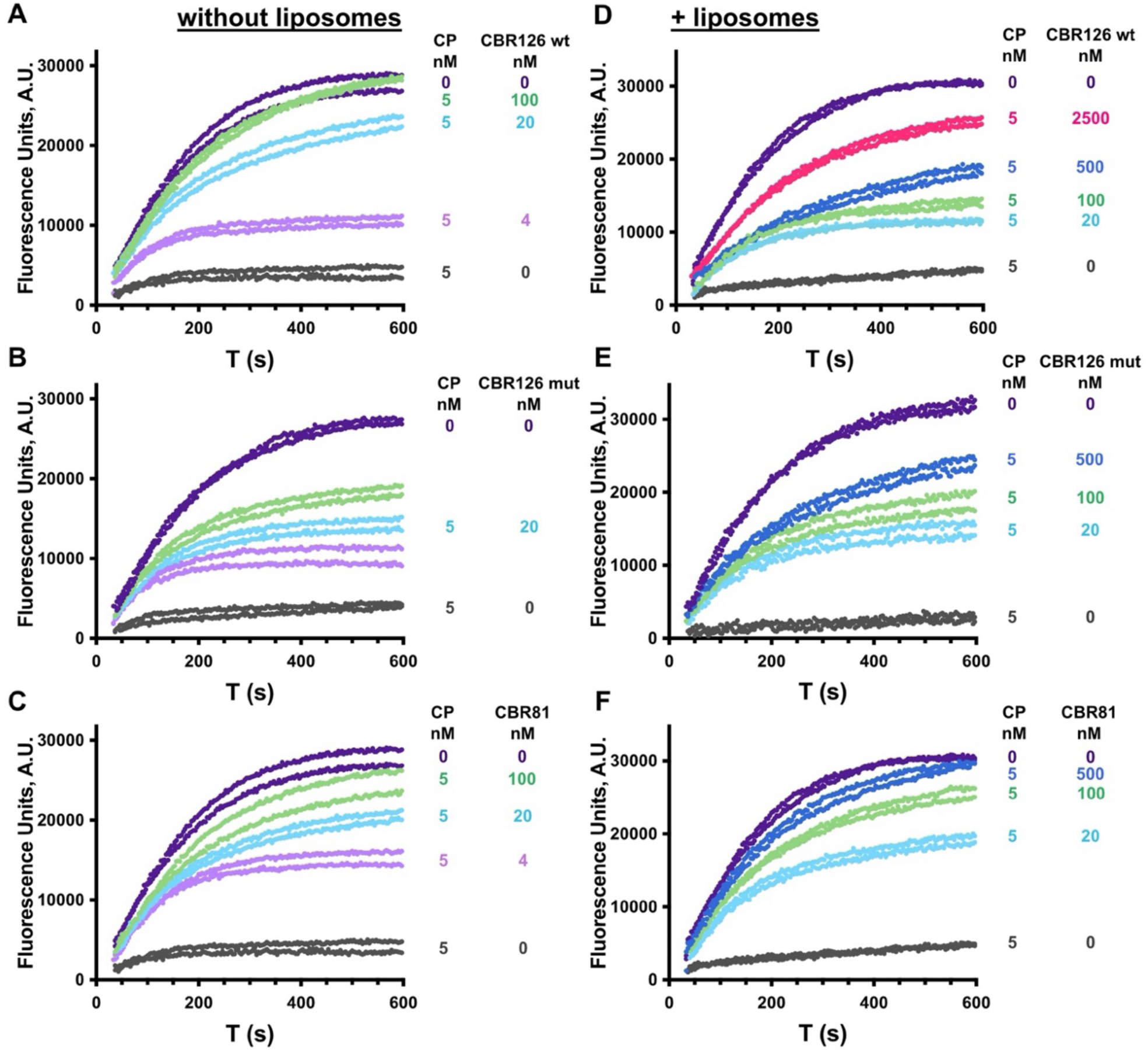
Actin polymerization capping assays – full set of CBR titration curves. Increasing concentrations of CBR126 wt (**A**), CBR126 mut (**B**), and CBR81 (**C**) were added to CP to inhibit its barbed-end actin capping activity. **D-F.** The same experiment as panels **A-C**, except in the presence of liposomes. Pyrene actin fluorescence (arbitrary units) is plotted vs time (s).

**Suppl. Figure 3.**
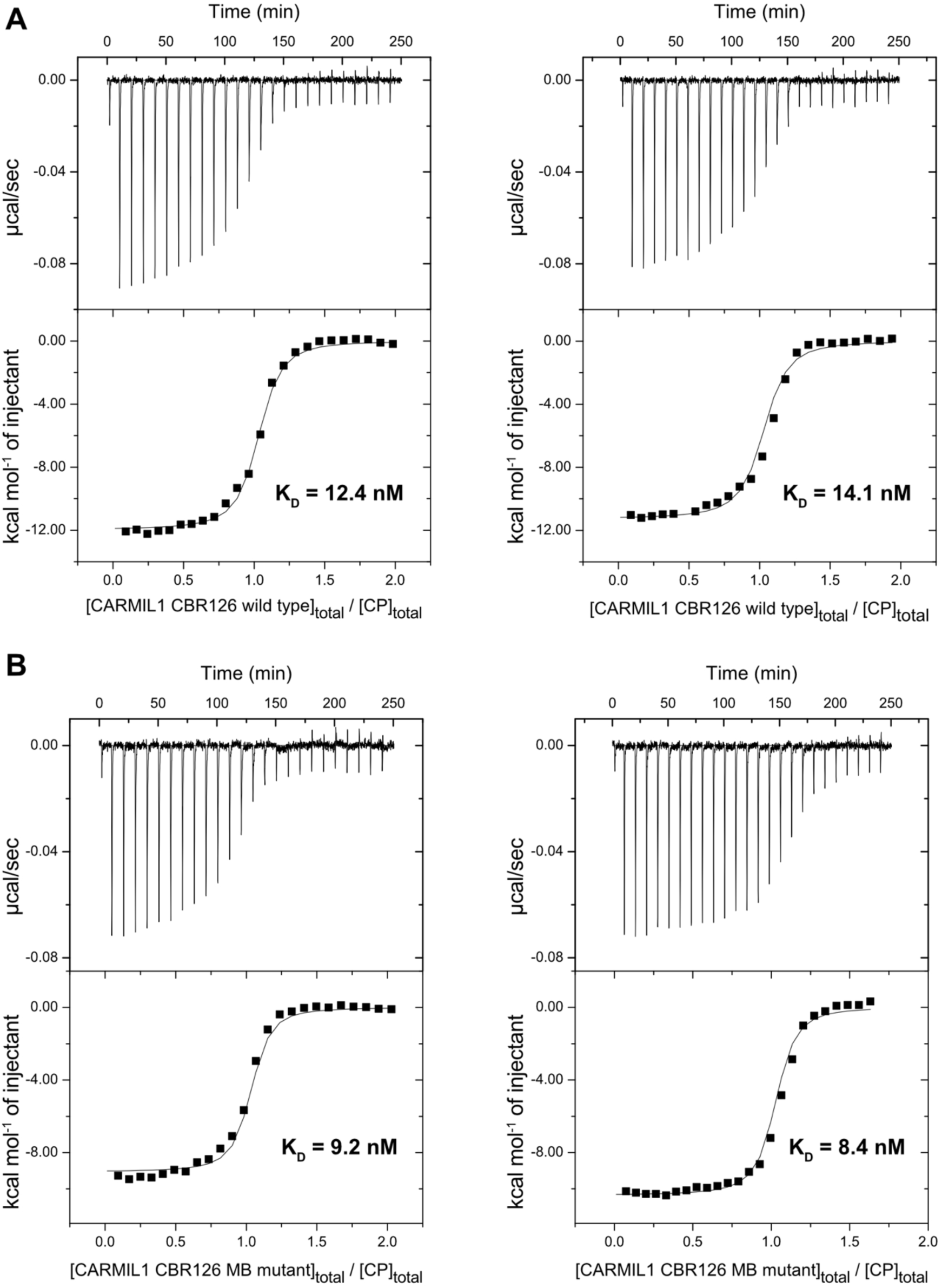
Binding affinities of CP for CBR126 wt and CBR126 mut are similar. ITC traces of CP titrated with CBR126 wt (panel A) or CBR126 mut (panel B). Upper panels are raw traces of thermograms, with differential power for each injection of CBR. Lower panels are binding isotherms, with integrated enthalpy vs molar ratio. Fitted values for K_D_, with a stoichiometry of one, are listed.

**Suppl. Figure 4.**
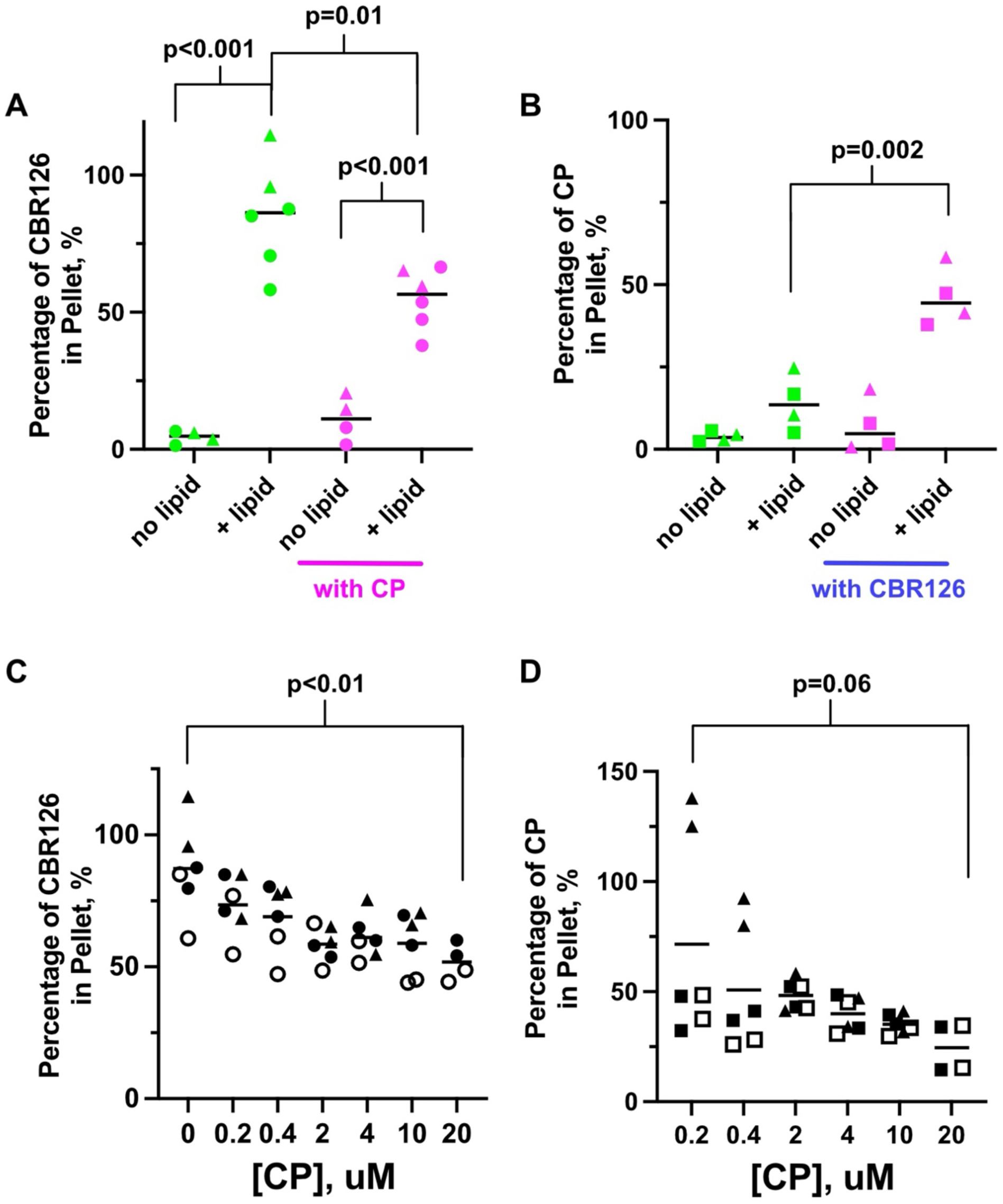
CP and liposomes appear to compete for CBR126 binding. **A.** Fraction of CBR126 that pellets with liposomes, plotted as a percentage of the total CBR126. CBR126 alone, without CP, sediments with liposomes when lipids are present, but not in the absence of lipids (green points). When CP is added to the reaction mixture, the amount of CBR126 that pellets with the liposomes is less (pink points). Each experiment is plotted as separate points, with the triangles representing a technical replicate set. The horizontal black bar is the median. P values from Welch’s t test analysis. **B.** Fraction of CP that pellets with liposomes, as a percentage of total CP. CP pellets with liposomes when CBR126 is present (pink points) but not in its absence (green points). Each experiment is plotted as separate points, with the triangles representing a technical replicate set. The horizontal black bar is the median. P values from Welch’s t test analysis. **C.** Fraction of CBR126 that pellets decreases as a function of increasing concentrations of CP. Experiments were performed in triplicate and technical replicate sets are shown as different shapes. Each value is plotted as a black horizontal line. P values from Welch’s t test analysis. **D.** Fraction of CP pelleting with CBR126 decreases with increasing concentrations of CP. The total amount of CBR126 was constant. Experiments were performed in triplicate and technical replicate sets are shown as different shapes. The horizontal black bar is the median. P values from Welch’s t test analysis.

**Suppl. Figure 5.**
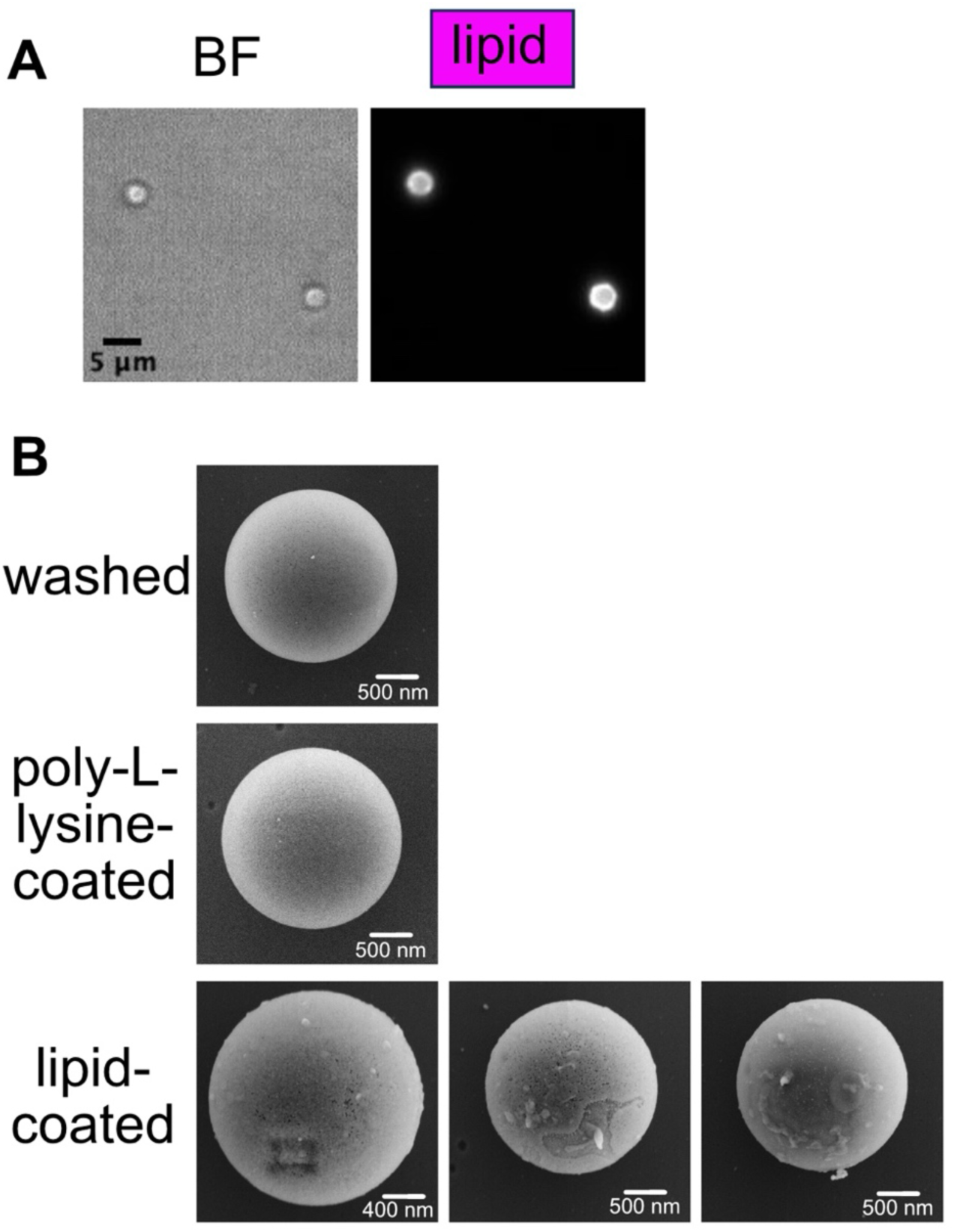
Lipid-coated silica beads. **A.** Brightfield (BF) and fluorescence image of beads coated with a lipid mixture containing 1% Cy5-PC. **B.** Scanning electron micrograph (SEM) images of silica beads either uncoated (washed), poly-L-lysine-coated, or lipid-coated.

**Suppl. Figure 6.**
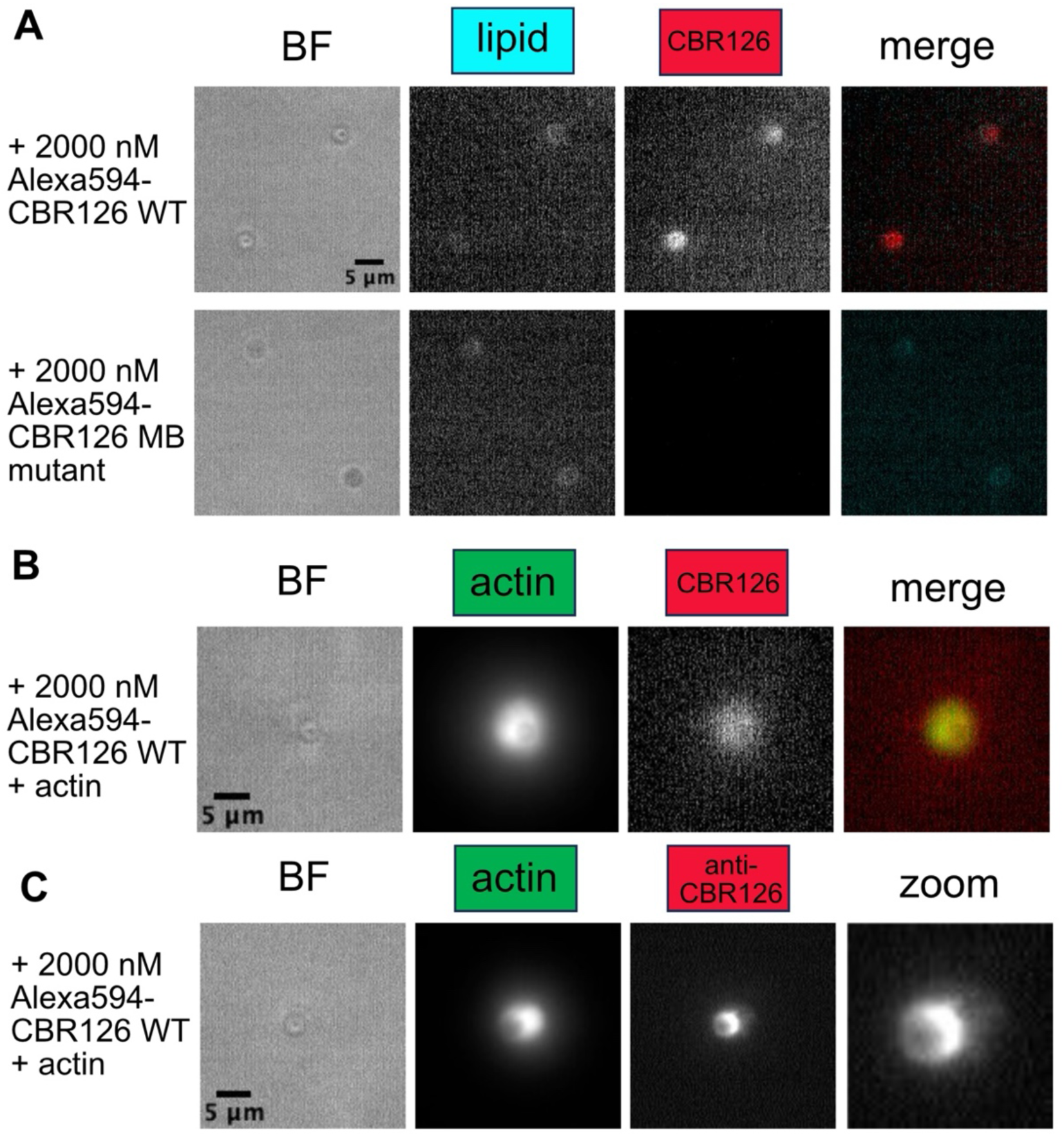
Localization of fluorescent CBR126 without His tag before and after actin polymerization. **A.** CBR126 wt localizes to the surface of the lipid-coated bead before addition of actin. CBR126 mut shows no localization. **B.** After actin polymerizes, CBR126 wt localization is lost at the bead surface, at times co-localizing with the F-actin. **C.** Localization of CBR126 by antibody staining of CBR126 confirms colocalization with actin. Image labeled “zoom” is a higher magnification of the anti-CBR126 image.

## Methods

Many of the Methods are similar to ones in a previous publication (12). Modifications to procedures are stated explicitly.

### Buffers

*Assay Buffer*: 20 mM HEPES, 100 mM KCl, 1 mM MgCl_2_, 1 mM EGTA, 1 mM TCEP, 1 mM ATP (pH 7.0). For bead actin polymerization fluorescence microscopy assays, the buffer included 0.2% methylcellulose (cP 400) to reduce Brownian motion, 2.5 mg/ml BSA to decrease non-specific sticking to the coverslip, and 20 mM beta-mercaptoethanol to lessen photobleaching.

### Proteins

*Actin, Arp2/3 Complex, Profilin, CP, V-1, and CARMIL1 CBR.* Unlabeled, pyrene-labeled, and Alexafluor 488-labeled gel-filtered G-actin from skeletal muscle α-actin (UniProt P68135, Oryctolagus cuniculus) were prepared as described (12). Porcine brain Arp2/3 complex from Cytoskeleton (Cat. No. RP01P, Denver, CO) was reconstituted per manufacturer instructions and used within one month. Human profilin-1 (UniProt P07737) was a gift from Dr. Silvia Jansen (Dept. of Cell Biology & Physiology, Washington University, St. Louis). Mouse CPα1β2 heterodimer (UniProt Q5RKN9 and Q923G3, pBJ 2041) and human V-1 (UniProt P58546, pBJ 2438) were purified as described (35).

CARMIL1 CBR126 wild type (pBJ 2491) was composed of MVKIH, a translational enhancement sequence (36), followed by HHHHH (His tag), LEVLFQGP (PreScission protease cleavage site), and human CARMIL1 (UniProt Q5VZK9) E964 to T1089. Codon-optimized DNA was synthesized and inserted into a pRSF-1b vector (Novagen, Madison, WI) by Azenta Life Sciences, Burlington, MA. Recombinant protein was expressed in E. coli NiCo21 (DE3) (New England Biolabs, Ipswich, MA). Cells were lysed, and the insoluble fraction of the lysate was dissolved in 20 mM sodium phosphate, 2 mM Tris(2-carboxyethyl)phosphine (TCEP), 1 mM NaN3, containing 6 M guanidine-HCl (pH 7.8). The protein was bound to Ni-NTA 6 Fast Flow (Cytiva, Marlborough, MA). Guanidine-HCl was washed from the Ni-NTA column with 20 mM Tris-HCl, 0.5 mM TCEP, 0.3 M NaCl, 1 mM NaN3 (pH 7.8), and the CARMIL peptide was eluted with 20 mM Tris-HCl, 0.5 mM TCEP, 0.3 M NaCl, 1 mM NaN3, 100 mM imidazole (pH 7.8). To prepare the tag-less CARMIL1 peptide, the 6x-His tagged peptide was cleaved with GST-tagged PreScission protease (Cytiva). The 6x-His tagged and tag-less peptides were each then purified by cation exchange chromatography using a POROS GoPure XS column (Applied Biosystems, Foster City, CA) in 20 mM sodium phosphate, 0.1 mM EDTA, 1 mM NaN3, 10 mM DTT (pH 7.5), eluted with a linear gradient to 1 M NaCl. Fractions containing purified peptide were pooled, dialyzed against 20 mM 3-(N-morpholino)propane-sulfonic acid (MOPS), 100 mM KCl, 1 mM TCEP, 1 mM NaN3 (pH 7.2), and stored at −70 C. Molar mass of the purified peptides was confirmed by MALDI mass spectrometry.

CARMIL1 CBR126 mut (pBJ 2549) preparation was similar to the CBR126 wt above (pBJ 2491) with several differences. The expressed construct was composed of MVKIH, a translational enhancement sequence (36), followed by HHHHH (His tag), LEVLFQGP (PreScission protease cleavage site), and a mutated version of human CARMIL1 (UniProt Q5VZK9) E964 to E1058 with a deletion of the BH motif. That mutated sequence was EEEEDSEESSGSSNSSESESESEEPPTISMT. Codon-optimized DNA was synthesized and inserted into a pRSF-1b vector (Novagen) by Azenta Life Sciences. Recombinant protein was expressed in E. coli NiCo21 (DE3) (New England Biolabs). Cells were lysed, and the insoluble fraction of the lysate was dissolved in 20 mM sodium phosphate, 2 mM TCEP, 1 mM NaN3, containing 6 M guanidine-HCl (pH 7.8). The protein was bound to Ni-NTA 6 Fast Flow (Cytiva). Guanidine-HCl was washed from the Ni-NTA column with 20 mM Tris-HCl, 0.5 mM TCEP, 0.3 M NaCl, 1 mM NaN3 (pH 7.8), and the CARMIL peptide was eluted with 20 mM Tris-HCl, 0.5 mM TCEP, 0.3 M NaCl, 1 mM NaN3, 100 mM imidazole, containing 6 M urea (pH 7.8). The CARMIL peptide was then purified by anion exchange chromatography using a POROS GoPure XQ column (Applied Biosystems) in 20 mM Tris-HCl, 1 mM NaN3, 10 mM DTT, containing 6 M urea (pH 7.5), eluted with a linear gradient to 1 M NaCl.

Fractions containing purified peptide were pooled and dialyzed against 20 mM MOPS, 100 mM KCl, 1 mM TCEP, 1 mM NaN3 (pH 7.2). To prepare the tag-less CARMIL1 peptide, the 6x-His tagged peptide was cleaved with glutathione-S-transferase (GST)-tagged PreScission protease (Cytiva) The GST-tagged PreScission protease was removed using Glutathione Superflow Agarose (Pierce Chemical Company, Dallas, TX). The peptides were stored at −70 C. Molar mass of the purified peptides was confirmed by MALDI mass spectrometry.

CARMIL1 CBR81 wild type (pBJ 2531) was composed of MVKIH, a translational enhancement sequence (36), followed by HHHHH (His tag), LEVLFQGP (PreScission protease cleavage site), and human CARMIL1 (UniProt Q5VZK9) E964 to T1044. Codon-optimized DNA was synthesized and inserted into a pRSF-1b vector (Novagen) by Azenta Life Sciences, Burlington, MA. Recombinant protein was expressed in E. coli NiCo21 (DE3) (New England Biolabs). Cells were lysed, and the insoluble fraction of the lysate was dissolved in 20 mM sodium phosphate, 2 mM Tris(2-carboxyethyl)phosphine (TCEP), 1 mM NaN3, containing 6 M guanidine-HCl (pH 7.8). The peptide was bound to Ni-NTA 6 Fast Flow (Cytiva). Guanidine-HCl was washed from the Ni-NTA with 20 mM Tris-HCl, 0.5 mM TCEP, 0.3 M NaCl, 1 mM NaN3 (pH 7.8) and the CARMIL peptide was eluted with 20 mM Tris-HCl, 0.5 mM TCEP, 0.3 M NaCl, 1 mM NaN3, 100 mM imidazole (pH 7.8). The protein was then purified by cation exchange chromatography using a POROS GoPure XS column (Applied Biosystems) in 20 mM sodium phosphate, 0.1 mM EDTA, 1 mM NaN3, 10 mM DTT (pH 7.5), eluted with a linear gradient to 1 M NaCl. Fractions containing purified peptide were pooled and dialyzed against 20 mM MOPS, 100 mM KCl, 1 mM TCEP, 1 mM NaN3 (pH 7.2). To prepare the tag-less CARMIL1 peptide, the 6x His tag was cleaved from the peptide with GST-tagged PreScission protease (Cytiva). The GST-tagged PreScission protease was removed using Glutathione Superflow Agarose (Pierce Chemical Company). The peptides were stored at −70 C. Molar mass of the purified peptides was confirmed by MALDI mass spectrometry.

#### AlexaFluor-594 Labeling of CBR126

Purified preparations of tag-less CBR126 wt and CBR126 mut were dialyzed against 20 mM sodium phosphate, 0.15 M NaCl, 1 mM NaN3 (pH 7.4). Six to ten mol of Alexafluor 594 C5 maleimide (Thermo Fisher Scientific) was added per mol of CBR, and the solution was incubated overnight at 4 C. The labeled CBRs were dialyzed against 20 mM sodium phosphate, 0.15 M NaCl, 1 mM NaN3, 10 mM DTT (pH 7.4), then 20 mM MOPS, 100 mM KCl, 1 mM TCEP, 1 mM NaN3 (pH 7.2), and stored at −70 C.

#### His-tagged VVCA

His-tagged VVCA (pBJ 2534) was composed of N-terminal GST-LEVLFQGP (PreScission protease cleavage site)-HHHHHH-human N-WASP (UniProt O00401) P392 to D505. Codon-optimized DNA was synthesized and inserted into a pRSF-1b vector (Novagen) by Azenta Life Sciences, Burlington, MA. The recombinant protein was expressed in E. coli NiCo21 (DE3) (New England Biolabs). The cells were lysed and the soluble fraction of the lysate was purified with Glutathione Sepharose 4 Fast Flow (Cytiva) in 20 mM sodium phosphate, 0.1 M NaCl, 0.1 mM EDTA, 1 mM NaN3, 10 mM DTT (pH 7.8), and eluted with 20 mM sodium phosphate, 0.1 M NaCl, 0.1 mM EDTA, 1 mM NaN3, 10 mM DTT, 10 mM reduced glutathione (pH 7.8). The protein was then purified by anion exchange chromatography using a POROS GoPure 50 HQ column (Applied Biosystems) in 20 mM Tris-HCl, 1 mM NaN3, 10 mM DTT (pH 7.5), eluted with a linear gradient to 1 M NaCl. Fractions containing purified peptide were pooled, dialyzed against 20 mM MOPS, 100 mM KCl, 1 mM TCEP, 1 mM NaN3 (pH 7.2), and stored at −70 C.

### Actin Polymerization Assays

Pyrene-labeled F-actin seeds were prepared by adding 1 mM MgCl2, 1 mM EGTA, 50 mM KCl, and 5 μM phalloidin to 5 μM G-actin (5% pyrene label) and incubating at room temperature overnight. To measure elongation rates, CP (0 or 5 nM) and tag-less CARMIL1 fragments (0-2500 nM), with or without 1 mg/mL 50% DOPC, 50% DOPS liposomes were then added to 2.4 µM G-actin (5% pyrene label). To initiate polymerization, pyrene-labeled F-actin seeds were added at a concentration of 1.25 µM. Elongation rates were measured at 25 °C using time-based scans on a steady state spectrofluorometer (PTI QuantaMaster 8000 with Felix GX software) with excitation at 368 nm and emission detected at 386 nm.

### Isothermal Calorimetry (ITC)

ITC experiments were performed on a MicroCal MicroCalorimeter (Malvern Panalytical, Malvern, PA). CP (2 μM) was titrated with tag-less CARMIL1 CBR126 wild type, tag-less CBR126 BH mutant, or tag-less CBR81 wild type (22 μM) at 25 °C in 20 mM MOPS, 0.3 M KCl, 1 mM TCEP, 1 mM NaN3, 0.005% Tween20 (pH 7.2). Binding constants were determined by fitting the change in enthalpy to a single site binding model using MicroCal ITC Origin analysis software.

### Liposome Preparation

#### Liposomes for Actin Polymerization Experiments

100 mg 18:1 (Δ9-Cis) PC (DOPC) [1,2-dioleoyl-sn-glycero-3-phosphocholine in chloroform (Avanti Research 850375C Alabaster, AL)] and 100 mg 18:1 PS (DOPS) [1,2-dioleoyl-sn-glycero-3-phospho-L-serine (sodium salt) in chloroform (Avanti Research 840035C)] were combined and dried with Argon gas followed by further drying under vacuum. 2 mL of 20 mM MOPS, 0.3 M KCl, 1 mM TCEP, 1 mM NaN3 (pH 7.2) was added to the dry lipids. The lipids were resuspended by vortexing and sonication. The lipid suspension was then frozen in a dry ice ethanol bath and thawed at room temperature. The freeze-thaw cycle was repeated four times. The lipid suspension was then extruded seven times through a 0.2 µm polycarbonate membrane (Avanti Research 610006). 1.8 mL of the liposome suspension was added to 7.2 mL 20 mM MOPS, 0.3 M KCl, 1 mM TCEP, 1 mM NaN3 (pH 7.2) and was stored at −70 C. Prior to use in assays, the liposome suspension was thawed and the extrusion was repeated.

#### Liposomes for Bead and Sedimentation Experiments

Lipids were mixed together to give a final total concentration of 5 mM at the following ratios: 44% 1,2-dioleoyl-sn-glycero-3-phosphocholine (18:1 (Δ9-Cis) PC, DOPC), Avanti Research, #850375, Alabaster, AL), 5% 1,2-dioleoyl-sn-glycero-3-[(N-(5-amino-1-carboxypentyl)iminodiacetic acid)succinyl] (nickel salt) (18:1 DGS-NTA(Ni), Avanti Research, #790404, Alabaster, AL), 50% 1,2-dioleoyl-sn-glycero-3-phospho-L-serine (sodium salt) (18:1 PS, DOPS, (Avanti Research, #840035, Alabaster, AL), 1% 1,2-dioleoyl-sn-glycero-3-phosphocholine-N-(Cyanine 5) (18:1 Cyanine 5 PC, Avanti Research, #850483, Alabaster, AL) or 1% 1,2-dioleoyl-sn-glycero-3-phosphoethanolamine-N-(1-pyrenesulfonyl) (ammonium salt) (18:1 pyrene PE, Avanti Research, #810331, Alabaster, AL). The lipid mixture was dried under argon, and then under vacuum for 1 hr. Lipids were re-suspended in MOPS buffer (20.0 mM MOPS, 0.1 M KCl, 1.0 mM NaN_3_, 1.0 mM TCEP, pH 7.2) and cycled through rounds of vortexing for 30 seconds and sonication for 5 minutes three times. Liposomes underwent 5 freeze/thaws using an ethanol/dry ice bath. Liposomes were then passed through a hand-extruder (Genizer, Avanti Research, #610000, Alabaster, AL) and through a 0.2 µm polycarbonate membrane (Avanti Research, #610006, Alabaster, AL) nine times and collected for use. Liposome preparation for sedimentation assays was the same except the lipids were 50% DOPC and 50% DOPS at a total final concentration of 20 mg/mL.

### Protein-Coating of Beads

Silica beads with a diameter of 2.5 µm (Bangs Laboratory, SS05000, Fishers, IN) were sonicated for 10 min, centrifuged (3 min, 5000 rpm), and washed using MOPS buffer (20.0 mM MOPS, 0.1 M KCl, 1.0 mM NaN_3_, 1.0 mM TCEP, pH 7.2). Beads were coated with poly-L-lysine (0.01% solution, Sigma, P4707, St. Louis, MO) by incubation for 1 hour at room temperature (r.t.). Poly-L-lysine-coated beads were then coated with lipid by mixing 3.5 µL of beads (8.0×10^6^ beads/µL, 6.2×10^6^ beads/µL final concentration) with 5 µL of 5 mM liposomes (preparation described above) in 45 µL MOPS buffer, sonicating three times for 10 seconds, and incubating for 1 hour at r.t. with frequent gentle agitation. Lipid-coated beads were washed twice in MOPS buffer. Beads were suspended in MOPS buffer containing the following concentrations of 6xHis-tagged fusion proteins: 2 µM His-VVCA, 0-2 µM His-CBR126, 0-2 µM Hexa His tag peptide (APExBIO, A6006, Houston, TX). Hexa His tag peptide was used as a placeholder when the concentration of His-CBR126 was less than 2 µM, so that the total concentration of 6xHis was 4 µM in all samples during the coating process. Beads were incubated with 6xHis fusion proteins for 2 hours on ice, with frequent agitation to maintain the beads in suspension. Beads were centrifuged and washed once in assay buffer (no ATP) with 1% BSA, followed by a second wash step in assay buffer (no ATP) with 0.1% BSA. Protein-coated beads were then suspended in assay buffer with 0.1% BSA and 1 mM ATP. Protein-coated beads were used within one or two days.

For assays that utilized tag-less CBR126, beads were first precoated with His-VVCA and Hexa His and washed as previously described. Next, these beads were incubated with various concentrations of tag-less fluorescently-labeled CBR126 wt or CBR126 mut for 15 min at r.t., followed by a centrifugation step and resuspension in assay buffer with 0.1% BSA and 1 mM ATP. We confirmed that CBR126 wt stayed associated with lipid beads over the course of several hours, but beads were always used immediately for the assays reported.

### Bead-based Actin Polymerization Microscopy Assays

For bead-based actin polymerization assays, protein-coated beads (∼3.2×10^5^ beads or ∼1.6×10^4^ beads / µL) were mixed with 100 nM Arp2/3 complex, CP (0-200 nM), 5 µM profilin, 5 µM actin (10% Alexa 488-labeled actin), and V-1 (0-5000 nM) in assay buffer containing 0.2% methylcellulose (cP 400), 2.5 mg/ml BSA, and 20 mM β-mercaptoethanol. To ensure that relevant complexes formed before the start of the actin polymerization reaction, we pre-mixed three pairs of components: Arp2/3 complex with VVCA-coated beads, actin monomers with profilin, and CP with V-1. The pairs were incubated for 15 min at RT. All three of the pre-mixes were then combined into one vessel to start the polymerization reaction. After 15 min at RT, the reaction mixture was mounted between a coverslip and a glass slide, sealed using clear fingernail polish and imaged after another 15 min. Alternatively, the reaction was arrested after 30 min of incubation by adding 10 µM phalloidin and 10 µM Latrunculin B, mounted between a coverslip and a glass slide, and imaged immediately. These reaction times were chosen after pilot experiments testing a range of times.

Samples were imaged using a 60x 1.45 NA oil objective on an inverted Olympus IX81 microscope using a mercury lamp for excitation. Images were collected in wide-field fluorescence and bright-field modes with a Hamamatsu EM-CCD (C9100) camera using Micro-Manager software (37). Digital images were analyzed using Fiji software (38).

### Antibody labeling of bead samples for imaging

Bead samples were prepared as described above, and reactions were arrested at 30-min with 10 µM phalloidin and 10 µM Latrunculin B (arrest mixture in assay buffer, no ATP). Beads were pelleted by centrifugation (10 min, 5000 rpm), suspended in arrest-mixture assay buffer with 3% BSA and 1:400 anti-CARMIL1 antibody (ThermoFisher, 27133-1-AP). After incubation for 2 hr at RT, beads were centrifuged and washed twice using arrest-mixture assay buffer. Secondary antibody (1:400 Alexa 647-labeled goat anti-rabbit Ig ThermoFisher A21244) was added in arrest-mixture assay buffer (no ATP) with 3% BSA. After incubation for 1 hour at RT, beads were washed twice, resuspended, and imaged as above.

### Scanning Electron Microscopy of Lipid-coated Bead Surfaces

Treated 2-µm silica bead samples (hereafter referred to as samples) were delivered in 217 mOsm/L MOPS buffer (3-(N-morpholino)propane sulfonic acid) and fixed in 0.1% osmium tetroxide for 30 minutes in Hanks Buffered Saline Solution (HBSS) adjusted to 200 mOsm/L. The samples were rinsed three times in the adjusted HBSS for 10 minutes each, then affixed to poly-L-lysine (1 mg/ml)-coated cover slips. The samples were dehydrated through a graded ethanol (ETOH) series (30%, 50%, 70%, 90%, and 100%) for 10 minutes at each concentration, repeated four times. The final two dehydrations in 100% ETOH were performed with ETOH dried over 3 Å molecular sieves (Sigma-Aldrich, St. Louis, MO). Once fully dehydrated, the samples were placed in a Leica EM CPD 300 critical point dryer (Vienna, Austria) set to perform 12 CO2 exchanges at the slowest speed. The sample coverslips were mounted on aluminum pin mounts with colloidal silver paint (Electron Microscopy Sciences, Hatfield, PA) and then coated with 10 nm of carbon and 6 nm of iridium using a Leica ACE 600 vacuum coater (Vienna, Austria). Scanning electron microscopy (SEM) images were captured using a Merlin (Carl Zeiss) field emission (FE)-SEM (Oberkochen, Germany) at a resolution of 2 nm at 3 kV and 200 pA probe current with a secondary electron detector.

## Abbreviations

CP: actin capping protein
F-actin: filamentous actin
CARMIL: capping protein, Arp2/3 and myosin I linker
CPI: capping-protein-interacting
CSI: CARMIL-specific-interacting
CBR: capping protein binding region
MB: membrane-binding domain
BH: basic-hydrophobic motif.

